# Hepatic stearoyl-CoA desaturase-1 is specifically suppressed by dextran sodium sulfate but does not influence colitis sensitivity

**DOI:** 10.64898/2026.05.29.728832

**Authors:** Camille Duchamp-Smith, Natalie Burchat, Lakshmi Gayathri Pantula, Samuel B. Mitchell, Tolunay B. Aydemir, Harini Sampath

## Abstract

The delta-9 desaturase stearoyl-CoA desaturase-1 (SCD1) catalyzes the conversion of saturated fatty acids to monounsaturated fatty acids (MUFA) and is highly expressed in liver and adipocytes. Previous studies have demonstrated that treating mice with dextran sulfate sodium (DSS), a chemical inducer of ulcerative colitis, results in severe downregulation of SCD1 in the liver. However, the specific role of hepatic SCD1 in modulating colitis severity, as well as the impact of DSS on SCD1 and other lipogenic factors in other tissues has not been investigated. Here we show that downregulation of hepatic SCD1 following DSS treatment is not accompanied by changes to other lipogenic genes in the liver. In contrast, adipose tissue demonstrates coordinated reductions in lipogenic genes, including SCD1 and SCD2, while the colon does not display any perturbation of these targets. Furthermore, we demonstrate that the downregulation of hepatic SCD1 occurs independently of sterol regulatory element binding protein-1c (SREBP-1c) and does not require an intact gut microbiome. Interestingly, a distinct model of colitis induced by IL-10 deficiency does not result in downregulation of hepatic SCD1. Concomitant with transcriptional changes, DSS treatment is associated with significant remodeling of the hepatic lipidome, including reductions in total phospholipids (PLs) and reduced MUFA-containing PLs and triacyglycerols (TAGs), consistent with the observed reduction in SCD1. Interestingly, hepatic cholesterol esters and plasma lipids including free cholesterol and glycerophospholipids were significantly elevated following DSS treatment. Given the significant reduction in hepatic SCD1 following DSS treatment, we tested a role for liver SCD1 in modulating colitis sensitivity. Mice with a targeted deletion of hepatic SCD1 were not more prone to colitis, indicating that the loss of hepatic SCD1, while a consequence of DSS-induced colitis, does not mediate colitis sensitivity in vivo.

**Synopsis:** Hepatic SCD1 does not modulate colitis severity upon DSS exposure. However, DSS-induced colitis elicits significant lipid metabolism dysfunction, demonstrated by elevated plasma and liver lipids, particularly plasma cholesterol and hepatic cholesterol esters, highlighting a role for gut-liver crosstalk following colonic inflammation.

## Introduction

Inflammatory bowel diseases (IBDs) including ulcerative colitis and Crohn’s disease are characterized by inflammation and injury to the small intestine and colon. Development of IBD involves multiple interconnected factors, including changes to gut permeability, genetic predispositions, unregulated immunological response, and host-bacteria interactions [1–4]. Beyond symptoms of gut inflammation, studies have also shown an increased prevalence of metabolic dysfunction-associated steatotic liver disease (MASLD), liver fibrosis and steatohepatitis, in patients with IBD compared to the general population [5, 6]. Metabolomic profiling of serum from IBD patients has demonstrated significant alterations in lipid- and TCA cycle-related metabolites compared to healthy subjects [7], suggesting that IBD is associated with systemic changes, beyond the gut. As the prevalence of IBD continues to rise in both industrialized and developing regions around the world [8], the need to understand IBD etiology and devise novel therapies that increase remission rates and delay or prevent secondary metabolic consequences are of paramount urgency.

Using both genetic and chemical means, multiple animal models of colonic inflammation and IBD have been studied [9–14]. A commonly used model includes the oral administration of the sulfated polysaccharide dextran sulfate sodium (DSS), which leads to colitis-like symptoms in mice when administered in drinking water [15]. Mice treated with DSS develop colonic crypt abscesses, mucosal immune cell infiltration, colon shortening, as well as watery and bloody fecal content. Beyond the gut, liver metabolomics studies have shown that metabolites associated with one-carbon metabolism and nucleotide synthesis are affected by acute DSS treatment in mice [16]. Chronic treatment with DSS results in severe dysregulation of lipid metabolism in liver and adipocytes, including reductions in lipogenesis and beta oxidation, as well as increased hepatic steatosis [17]. However, alterations in hepatic lipid metabolism in commonly used short-term DSS-colitis models have not been thoroughly investigated.

Stearoyl-CoA desaturase (SCD) is a lipogenic enzyme that catalyzes the conversion of saturated fatty acids (SFAs) to monounsaturated fatty acids (MUFAs) [18]. SCD1 is ubiquitously expressed in the mouse, with high expression in adult liver, where it is induced by nutritional cues such as high-carbohydrate feeding and suppressed by fasting or feeding of polyunsaturated fatty acids [19–22]. Both acute and chronic DSS models have been reported to reduce hepatic SCD1 [17, 23]. Acute DSS reduces hepatic SCD1 in a dose and time dependent manner without any apparent impacts to other lipogenic genes, and also elicits reductions in plasma MUFAs, particularly oleoyl-lysophosphatidlycholine (18:1-LPC) [23]. Meanwhile, chronic DSS treatment reduces hepatic lipogenesis, including, but not limited to SCD1 expression[17]. Mice with whole-body deletion of SCD1 (*Scd1*^−/−^) were administered DSS in their drinking water and shown to be more prone to colitis than their WT controls [23]. Conversely, adenovirus-mediated overexpression of SCD1, thought to mostly result in elevated hepatic expression, was reported to ameliorate colitis symptoms in WT mice, suggesting that hepatic SCD1 may modulate colitis risk [23]. However, the impact of liver-specific SCD1 loss, with intact expression in other tissues such as the gut where the impact of DSS is most pronounced, has not been examined.

In the current study, we investigated the impact of acute oral DSS on expression of SCD1 and other lipogenic genes in the liver, adipose tissue, and colon. In addition, we investigated lipidomic changes in liver and plasma in response to acute DSS treatment. Further, we utilized a model of liver-specific deletion of SCD1 (LKO mice) to assess the distinct role of hepatic SCD1 in modulating the response to oral DSS.

## Results

### Acute DSS exposure lowers hepatic SCD1 without altering other lipogenic genes

Acute DSS treatment has been reported to sharply reduce hepatic SCD1 expression [23], but its impact on other hepatic lipogenic genes or lipid metabolism is not known. To investigate the impact of acute DSS treatment on hepatic lipogenic genes and lipid content, wild type (WT) male and female mice were given 3% DSS in drinking water for 5 days. Colitis was successfully established as evidenced by significant body weight loss (Figs. 1A and 1C) and appearance of occult blood in the DSS-treated groups by day 5 of DSS exposure (Figs.1B and 1D). Histological analyses of colon tissues showed loss of crypt structure with increased immune cell infiltration in the DSS-treated group, compared to the control group (Fig. 1E). Interestingly, despite no differences in food intake (Fig. 1F), both males and females on DSS showed increase water intake (Fig. 1G). Hepatic SCD1 was severely reduced at both the gene and protein levels following DSS treatment, in both male and female mice (Figs. 1H-I). Since responses were similar between sexes, the remainder of the study predominantly studied male mice, unless specifically noted.

**Figure 1:**
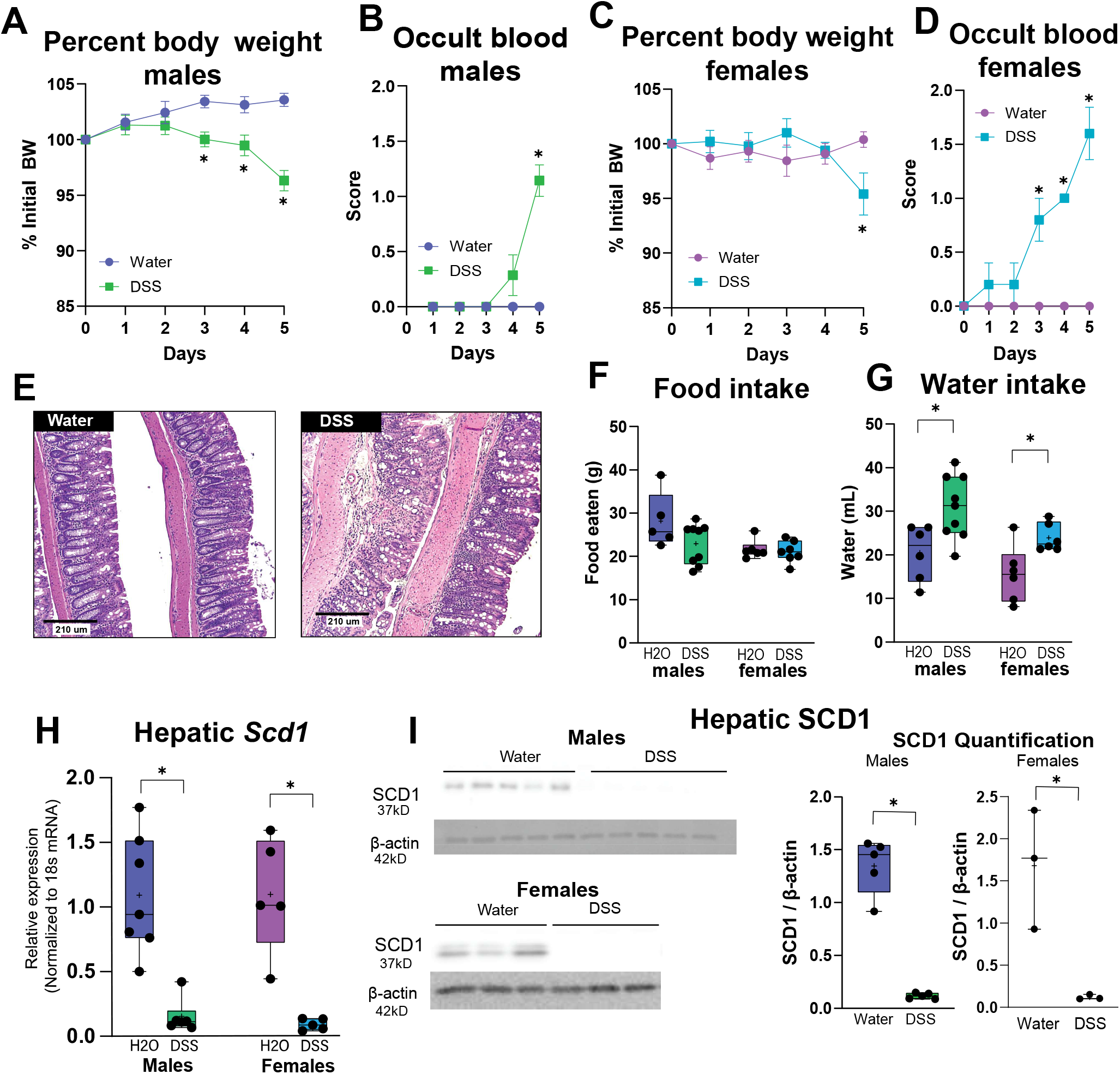
Oral DSS causes colitis and reduces hepatic SCD1 expression. (A, C) Bodyweights and (B, D) occult blood scores for mice given water (control) or 3% DSS in drinking water. Occult blood was scored as follows: 0, not occult blood; 1, positive occult blood; 2 visible blood in feces; 3, bloody anus. (E) H&E stain of colons from male control mice (left) or 3% DSS-treated mice (right). Scale bars represent 210 μm (F) Food intake and (G) water intake measured over the course of the study. (H) *Scd1* gene expression and (I) SCD1 protein expression in livers of control and DSS-treated mice. n=3-8 per group. Box plots show minimum and maximum values, + symbol at mean. *p<0.05 vs. water group.

Interestingly, expression of sterol regulatory element binding protein-1C *(Srebp*-1c), a key transcriptional regulator of *Scd1* and other lipogenic genes, was not altered by DSS (Fig 2A). Consistent with this lack of regulation of *Srebp-1c*, expression of other *Srebp-1c* targets, including acetyl-CoA carboxylase *(Acc)* and fatty acid synthase *(Fas*) was unchanged in livers of DSS-treated mice (Fig. 2A). Expression of the diacylglycerol O-acyltransferase −1 and −2 (*Dgat1* and *Dgat2)* isoforms was slightly reduced in livers of DSS-treated mice (Fig. 2A). Genes regulating hepatic lipid oxidation were also not impacted by DSS treatment (Fig. 2B). A significant reduction in expression of the fatty acid transporter *Cd36* suggested a possible reduction in hepatic lipid uptake (Fig. 2B). Taken together, these data indicate that DSS induces specific reductions in hepatic SCD1, without altering other lipid metabolic genes, likely via an SREPB-1c-independent mechanism.

**Figure 2:**
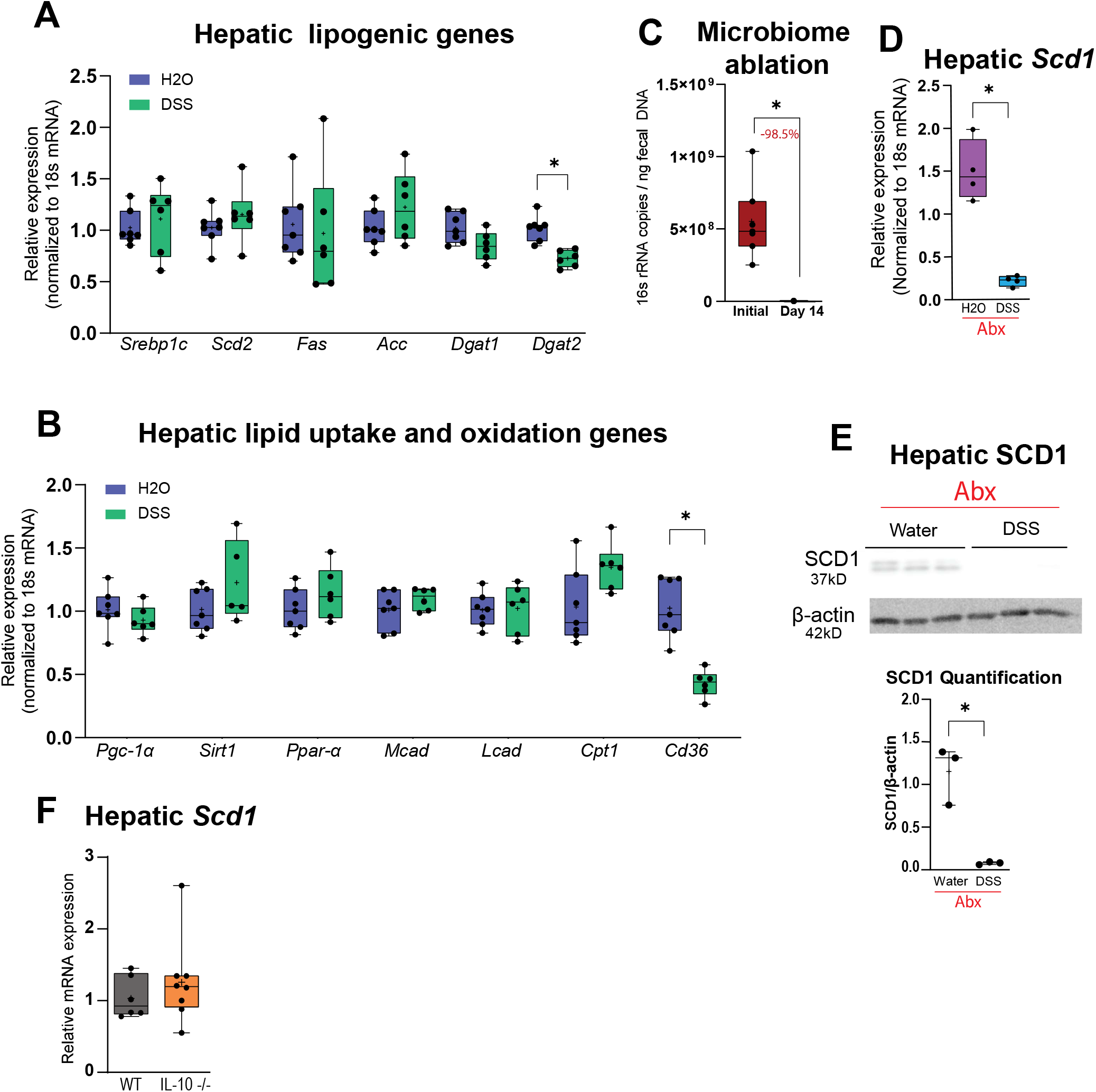
Specificity of and mechanisms underlying modulation of hepatic SCD1 by oral DSS. Expression of (A) lipogenic genes and (B) lipid uptake and oxidation genes were measured via qRT-PCR in liver of water and DSS-treated male mice. (C) Confirmation of microbiome ablation on day 14 following the start of Abx via qPCR of fecal 16s rRNA in female mice. Liver (D) gene and (E) protein expression of SCD1 following Abx treatment. (F) *Scd1* expression in livers of *IL-10*^−/−^ mice measured by qRT-PCR. n=3-8 per group. Box plots show minimum and maximum values, + symbol at mean. *p<0.05 vs. water group.

Treatment of mice with DSS is known to severely affect gut permeability via loss of the tight junction protein ZO-1 [24], leading to leakage of luminal content, including bacterial populations, across the epithelial barrier and triggering an inflammatory response [25]. To investigate whether the signal responsible for DSS-induced hepatic SCD1 reduction could be of microbial origin following acute damage to the colonic epithelium, we treated WT mice with an antibiotic (abx) cocktail for two weeks to ablate their gut microbiome. Fecal pellets were collected prior to abx start and after 2 weeks, to confirm successful microbiome ablation via fecal 18s rRNA expression (Fig. 2C). Abx-treated mice were subjected to oral DSS (5 days, 3% DSS). Interestingly, the reduction in hepatic SCD1 following acute DSS treatment was maintained in abx-treated mice (Figs. 2D, E), suggesting a microbiome independent mechanism for the observed reduction in hepatic SCD1. Further, to test whether hepatic SCD1 is generally suppressed in a distinct model of chronic IBD, we measured hepatic SCD1 expression in livers of mice lacking interleukin-10 (*IL-10*^−/−^) mice [14]. Interestingly, *IL-10*^−/−^ mice did not display a similar reduction in hepatic *Scd1* (Fig. 2F), speaking to important differences between acute DSS induced colitis vs. chronic immune-mediated IBD present in the *IL-10*^−/−^ model.

### Acute DSS treatment remodels the hepatic lipidome

Given the role of SCD1 in converting SFAs to MUFAs, we investigated whether the acute reduction in hepatic SCD1 following DSS treatment led to changes in hepatic lipid content and composition. Total hepatic lipids were significantly reduced by 17% following DSS treatment (Fig. 3A). This included a significant 11% reduction in hepatic phospholipids (PLs) (Fig. 3B), driven by reductions in MUFA products of SCD1, palmitoleate (C16:1) and oleate (C18:1) as well as several polyunsaturated fatty acids (PUFAs) within the PL fraction (Fig. 3E). As a percentage of liver PLs, DSS-treatment increased the proportion of SFAs and reduced MUFA-PLs (Fig. 3H), consistent with the observed reduction in hepatic SCD1 (Figs. 1H, I). Alterations in PUFA to SFA ratios in the PLs, a predominant constituent of cell membranes, can lead to changes in membrane fluidity and trigger processes like ER stress [26, 27]. However, no differences in markers of ER stress, including spliced XBP-1, *Gadd34*, or *Atf6* were observed following this acute DSS treatment (Fig 3I). No differences were noted in levels of total hepatic triacylglycerols (TAG) between control and DSS-treated mice (Fig. 3B). However, DSS treatment significantly reduced TAG myristoleate (C14:1) levels, and led to non-significant reductions in other SCD1 products, including C16:1 and C18:1 in the TAG fraction (Fig. 3F). Additionally, the PUFAs linolenic acid (C18:2n6) and gamma linolenic acid (C18:3n6) were significantly increased in the TAG fraction (Fig. 3F). As a result, despite a lack of significant changes in total amounts, the percent composition for hepatic TAGs indicated a significant reduction in SFAs and MUFAs and a significant increase in PUFAs (Fig. 3H) in DSS-treated mice. In addition to lipid acyl chain composition, five desaturation ratios, namely 14:1n5/14:0, 16:1n7/16:0, 18:1n7/18:0, and 18:1n9/18:0, as well as a total desaturation ratio indicating the sum of MUFAs/ sum of SFAs, were calculated for each lipid fraction to assess the impact of reduced hepatic SCD1 levels. These analyses revealed a significant reduction in total desaturation ratios in PLs and TAGs in DSS-treated livers, reflected as significant reductions in various desaturation indices in both lipid fractions (Fig. 3J). Together, these results indicate that the reduction in hepatic SCD1 following DSS treatment alters the hepatic lipidome by reducing MUFA incorporation into PLs and TAGs.

**Figure 3.**
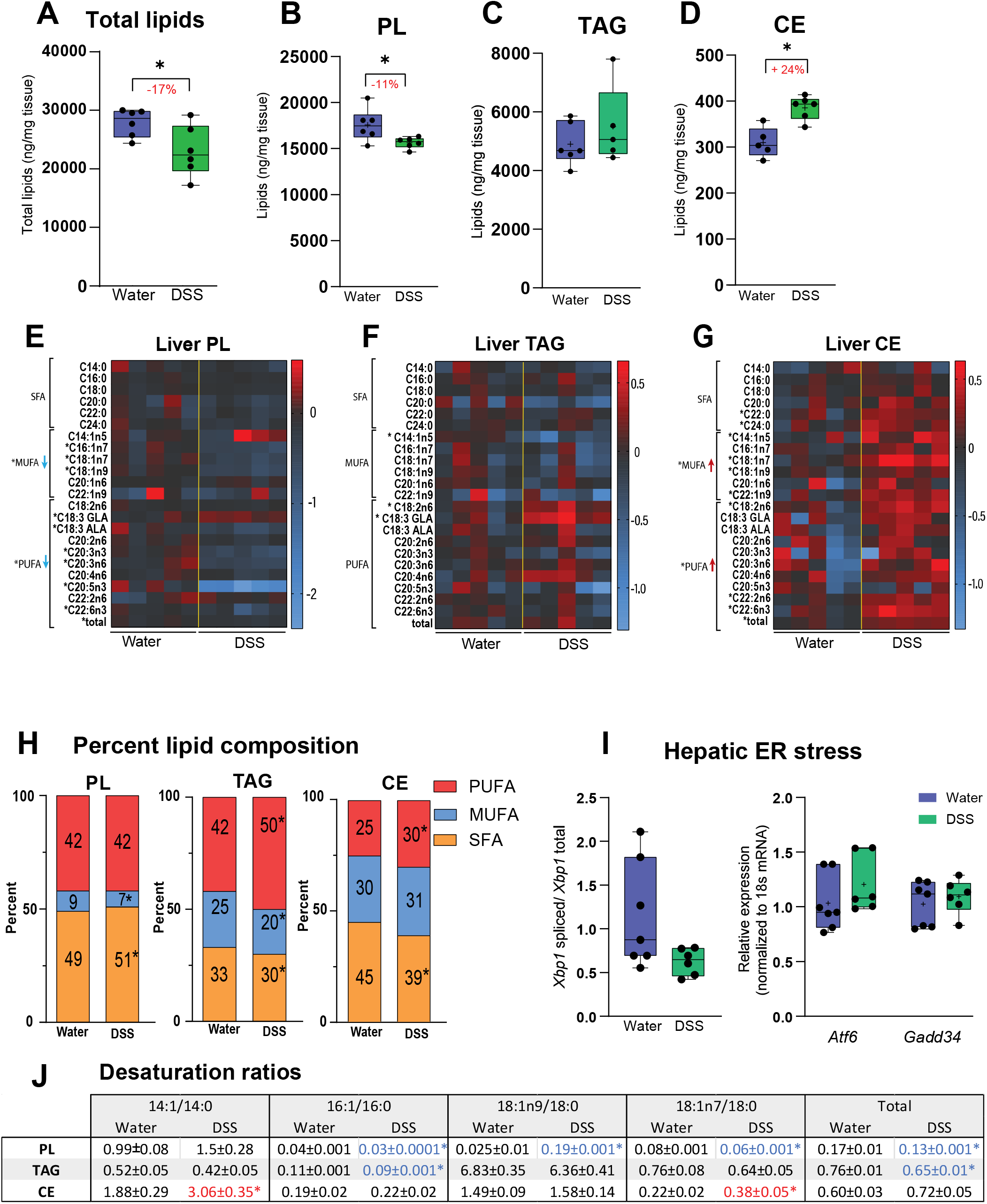
(Above): Acute DSS triggers changes in liver lipid composition. (A-G) Total liver lipid content, including total liver phospholipid (PL), triacylglycerol (TAG) and cholesterol ester (CE) content and composition. (H) Percent composition of saturated fatty acids (SFA), monounsaturated fatty acids (MUFA) and polyunsaturated fatty acids (PUFA) in PL, TAG and CE fractions. (I) Hepatic gene of ER stress markers measured via qRT-PCR. (J) Desaturation ratios for 14-18 carbon chain lipid species were calculated for each lipid fraction (PL, TAG, CE). Blue text denotes desaturation ratios that were significantly decreased, and red text denotes desaturation ratios that were significantly increased in the DSS-treated group, compared to the water group. n=5-7 per group. Box plots show minimum and maximum values, + symbol at mean. *p<0.05 vs. water group.

Apart from these changes, we noted a striking 24% increase in hepatic cholesterol esters (CEs) following DSS treatment (Fig 3D). Interestingly, this increase was driven mainly by a significant increase in MUFAs (C14:1, C18:1 and C22:1), as well as the PUFA C18:2n6 (Fig. 3G). As a relative percentage, hepatic CEs from DSS-treated mice had significantly higher %PUFAs and lower %SFAs (Fig. 3H). Interestingly, individual desaturation ratios were significantly elevated in the CE fraction, in contrast to hepatic PLs and TAGs (Fig. 3I). Together, these data indicate preferential channeling of MUFAs to hepatic CEs in DSS-treated mice, thus increasing CE-MUFA content despite the striking reduction in SCD1 expression. Other lipid species measured, including diacylglycerols, monoacylglycerols, and free fatty acids, did not show any major compositional changes following DSS treatment (Supplementary Figs. 3A-F).

### Hepatic cholesterol metabolism is altered upon DSS treatment

Given the striking changes in CE content in livers of DSS-treated mice, we measured expression of genes regulating cholesterol metabolism. Expression of 3-hydroxy-3-methylglutaryl-coenzyme A reductase (*Hmgcr)*, that catalyzes the rate-determining reaction for cholesterol biosynthesis in the liver, was downregulated in DSS-treated mice. Genes regulating hepatic cholesterol export via cholesterol transporter ATP-binding cassette transporter A1 *(Abca1)* and microsomal triglyceride transfer protein *(Mttp)*, facilitating cholesterol efflux into lipoproteins, were both significantly increased (Fig. 4A). Additionally, expression of scavenger receptor class B type 1 (*Scarb1)* (Fig. 4B) was also elevated, suggesting a potential increase in cholesterol uptake via high-density lipoprotein (HDL). Conversely, expression of the low-density lipoprotein receptor *(Ldlr)* was significantly reduced (Fig. 4B). Expression of the acyl-CoA:cholesterol acyltransferases 1 and 2 (*Acat1 and Acat2)*, which catalyze the cholesterol esterification process, was unchanged after DSS treatment (Fig. 4C). A previous study reported significant reductions in hepatic bile acid synthesis upon DSS treatment [28]. Consistently, we also observed that expression of genes encoding critical enzymes *Cyp7A1* and *Cyp27A1* was significantly downregulated in DSS-treated mice (Fig. 4C). Additionally, bile acid transporters *Ntcp, Oatp1, Mrp2*, and *Mrp3* were also significantly decreased (Fig. 4D), with a notable exception for Ost-β, which was upregulated (Fig. 4D) and *Bsep* which was not changed (Fig. 4D) by DSS treatment. Expression of the key transcriptional regulator of bile acid metabolism, farnesoid X receptor (*Fxr)* was unchanged. However, its target and known suppressor of bile acid synthetic pathways, small heterodimer partner (Shp), was significantly upregulated following DSS treatment (Fig. 4E), consistent with the observed reductions in *Cyp7a1*.

**Figure 4:**
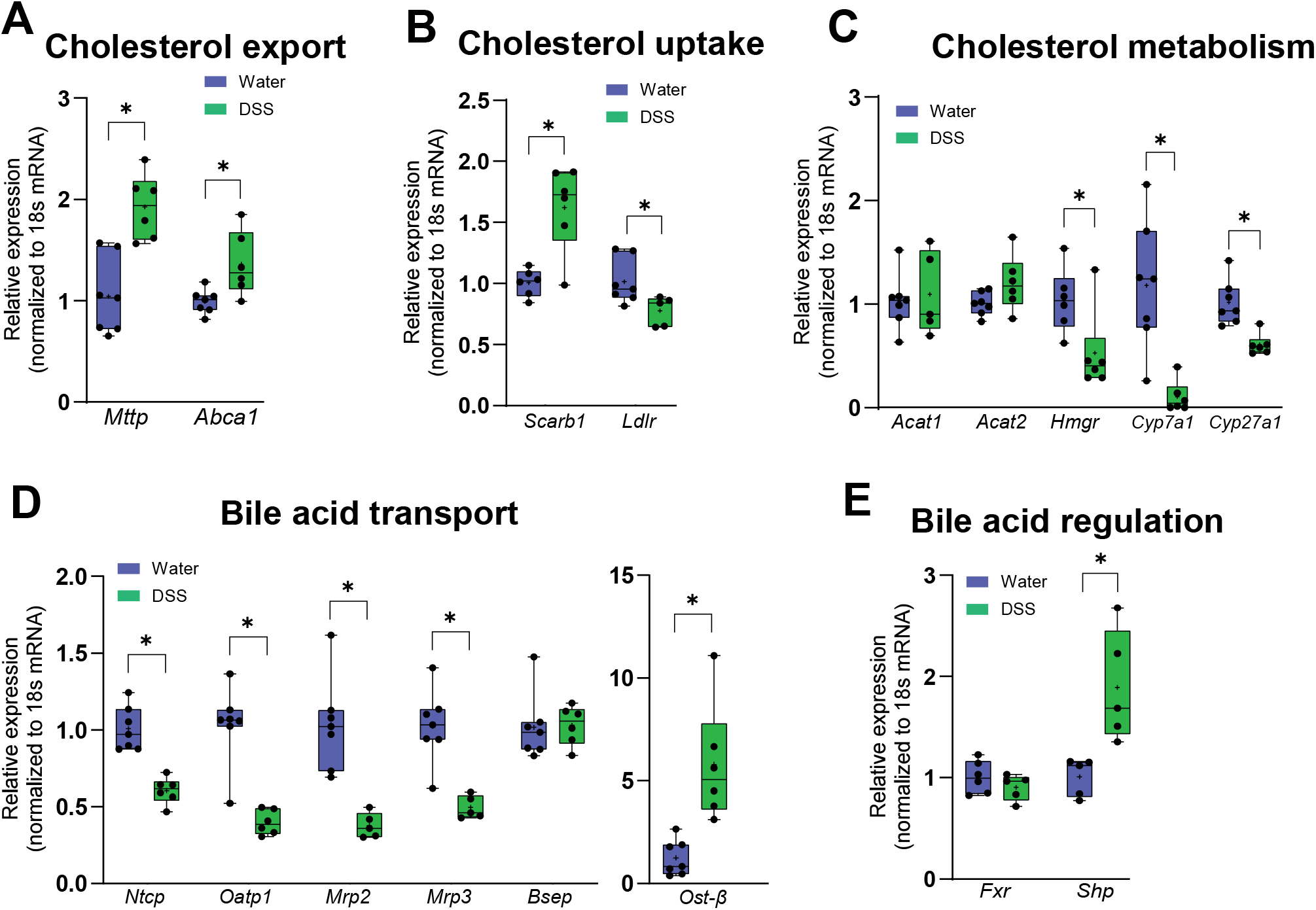
Hepatic cholesterol metabolism is affected following acute DSS. (A-E) Hepatic gene expression measured via qRT-PCR. n=5-7 per group. Box plots show minimum and maximum values, + symbol at mean. *p<0.05 vs. water group.

These data indicate that increased content of hepatic CEs is not driven by increases in cholesterol biosynthesis or esterification, but rather from an almost complete (90%) reduction in BA synthesis gene *Cyp7a1* via *Shp* signaling, leading to increased cholesterol buildup and export out of the liver into circulation.

### Acute DSS triggers increase in plasma lipids

Total plasma lipids were significantly elevated in DSS-treated animals (Fig. 5A), driven by significant increases in SFAs and PUFAs, without any changes noted in absolute MUFA levels (Fig. 5B). The relative percentage of PUFAs was elevated in plasma of DSS-treated mice, while percent MUFAs was significantly reduced (Fig. 5C). Further analysis of plasma lipids via LC-MS indicated several significant changes. Free cholesterol (FC) was strikingly increased by 60% in plasma of DSS-treated mice, without any changes noted in levels of plasma CEs (Fig. 5D). This increase in plasma cholesterol was also confirmed by colorimetric assay (Fig. 5E). To understand the localization of the increased plasma cholesterol, plasma lipoproteins were fractionated, and cholesterol associated with the VLDL, LDL, and HDL fractions were quantified in plasma from control and DSS-treated mice. Cholesterol was not detectable in very-low density lipoproteins (VLDLs) as expected, while cholesterol content was increased in both LDL and HDL fractions of DSS-treated mice (Fig. 5E). This elevation in plasma cholesterol is likely a consequence of acute and significant reduction in bile acid synthesis in DSS-treated mice (Fig. 5C).

**Figure 5.**
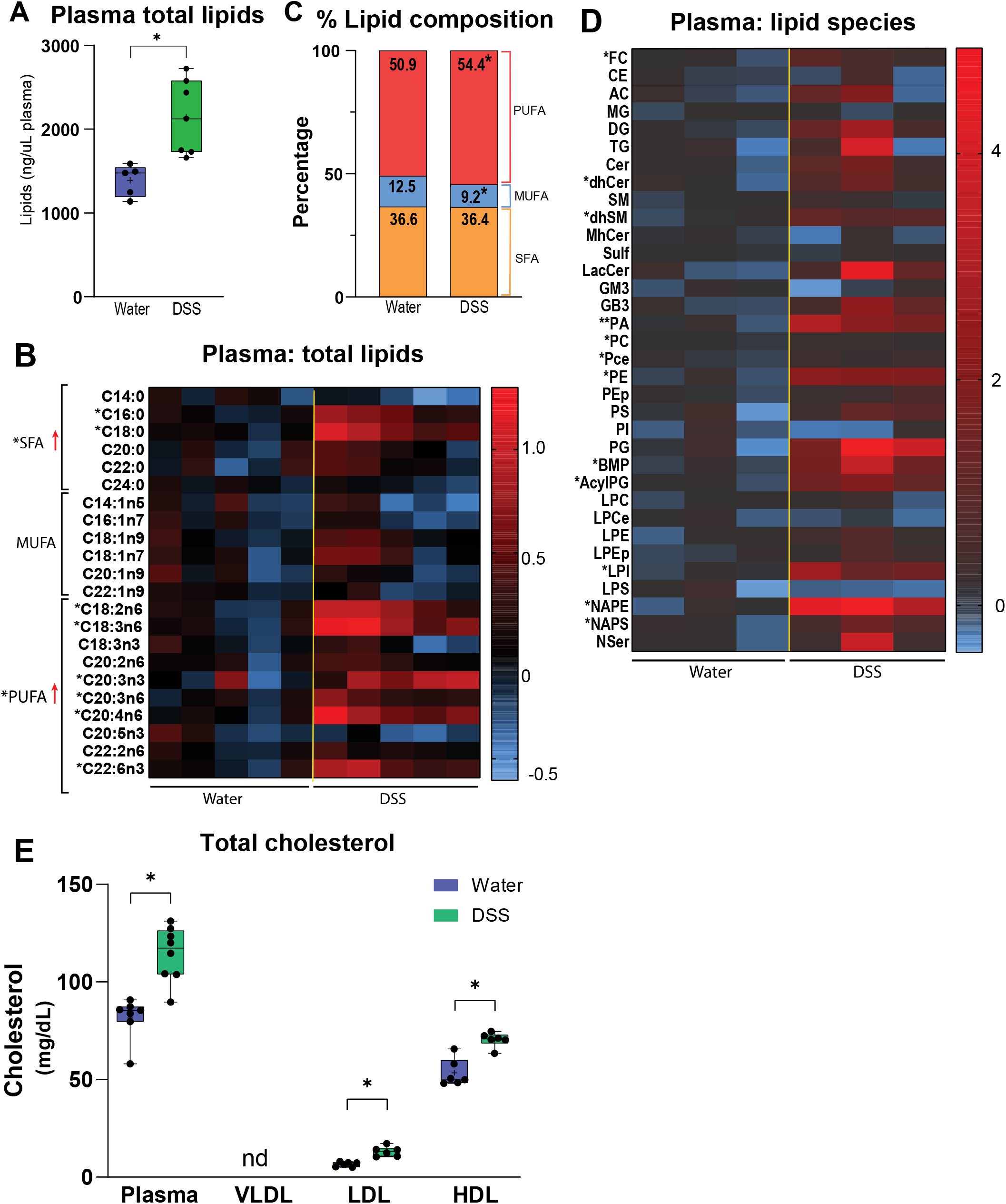
(Above): Acute DSS triggers increase in plasma lipids. (A-B) Total plasma lipids measured by GC-MS, n=5-7 per group. (C) Percent composition for total lipids was calculated. (D) Plasma lipidomic profiling was performed via UPLC-MS/MS n=3 per group. (E) Plasma cholesterol measured via colorimetric assay, n=7-8 per group. Measurement of total cholesterol in VLDL (not detected, nd), LDL and HDL performed following lipoprotein fractionation of plasma, n=6 per group. Box plots show minimum and maximum values, + symbol at mean. *p<0.05 vs water group.

In addition to changes in free cholesterol, several PL species were elevated in plasma of DSS-treated mice, including phosphatidic acid (PA), phosphatidylcholine (PC), alkyl phosphatidylcholine (PCe), phosphatidylethanolamine (PE), bis(monoacylglycero)phosphate (BMP or LBPA), (AcylPG) acyl phosphatidylglycerol, lysophosphatidylinositol (LPI), N-acyl phosphatidylethanolamine (NAPE) and N-Acyl phosphatidylserine (NAPS) (Fig. 5D). The increase in total levels of these PL species was reflected by increased levels of various acyl chains, both saturated and unsaturated, within each fraction (Supplementary Fig. 2). Increased plasma PLs may also be functionally related to reductions in bile acid synthesis in DSS-treated mice or be driven by a requirement for PLs in the colon to support recovery from injury.

### Altered lipid metabolism in colon and eWAT following acute DSS treatment

To investigate whether the severe reduction in hepatic SCD1 following acute DSS treatment was limited to the liver or could be observed in other tissues, SCD1 levels were assessed in the colon, the main site of DSS-induced damage and in epididymal white adipose tissue (eWAT), a site of high SCD1 expression. Following DSS treatment, colonic SCD1 expression remained unchanged (Fig. 6A). However, a moderate but significant reduction in *Srebp-1c* and *Acc* expression was observed in the colon following DSS treatment (Fig. 6C). In eWAT, SCD1 expression was reduced (Figs. 6B, D), similar to hepatic downregulation. In contrast to liver, SCD1 downregulation in WAT was accompanied by concomitant downregulation of other lipogenic genes, including *Srebp1c* and its target genes *Fas* and *Acc* (Fig. 6D). This coordinated reduction of lipogenic genes suggested an overall reduction in anabolic markers following DSS treatment rather than a specific regulation of SCD1 as seen in the liver. This was further supported by increased expression of adipose triglyceride lipase *(Atgl)* (Fig. 6E), suggesting increased lipolysis.

**Figure 6:**
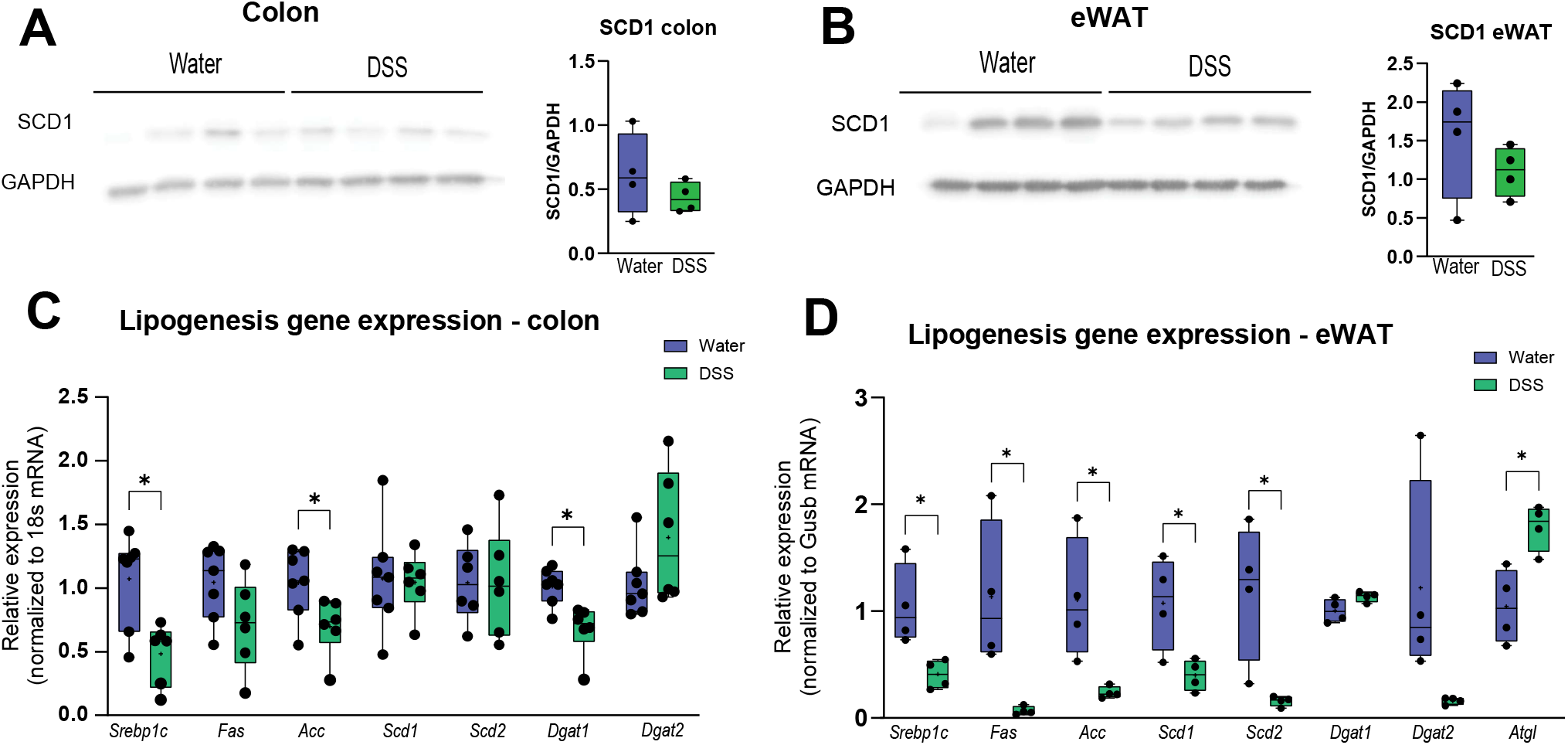
Lipid metabolism changes in colon and eWAT following Acute DSS. SCD1 protein expression in (A) colon and (B) eWAT of water and DSS-treated mice determined by western blot. Expression of lipogenic genes in (C) colon and (D) eWAT was measured via qRT-PCR in control and DSS groups. n= 4-6 per group. Box plots show minimum and maximum values, + symbol at mean. *p<0.05 vs water group.

### Loss of hepatic SCD1 does not sensitize mice to DSS-colitis

A previous report suggested that hepatic SCD1 may mediate the proinflammatory response to DSS [23]. However, this study was carried out in mice lacking whole-body SCD1 (*Scd1*^−/−^), thus making it difficult to deconvolute the role of hepatic SCD1 from its role in other tissues. The impact of specific loss of hepatic SCD1 on sensitivity to colitis has only been presumed but never directly tested. To test a causative role for hepatic SCD1 in DSS-induced colitis, mice lacking liver SCD1 expression were generated. Briefly, mice expressing Cre recombinase under the albumin promoter were crossed with mice expressing LoxP surrounding the *Scd1* gene. Mice expressing Cre recombinase served as liver SCD1 knockout mice (LKO), while floxed littermates not expressing *Cre* served as control mice (Flox) (Fig. 7A). Hepatic SCD1 expression was ablated in these mice (Fig. 7B), and the efficacy and specificity of this model have also been previously reported [20]. Flox and LKO littermates were given 3% DSS in drinking water for 5 days. Unlike whole-body *Scd1*^−/−^ mice that were previously studied [23], LKO mice drank the same amount of water as flox controls during DSS treatment (Fig. 7C). Daily monitoring showed no differences in body weight loss (Fig. 7D) or occult blood scores (Fig. 7E) between flox and LKO. Colon lengths were comparable between both groups (Fig. 7F), indicating no increase in colonic damage due to loss of hepatic SCD1. Further, colonic expression of inflammatory cytokines *Lcn2, Il-1b*, and *Il-6* were not significantly increased in LKO mice treated with DSS (Fig. 7G). Plasma levels of lipocalin-2 (LCN2), a marker of colonic inflammation, were not significantly increased in LKO animals (Fig. 7H). Taken together, these results indicate that while suppression of hepatic SCD1 is a consequence of acute DSS exposure, loss of hepatic SCD1 alone does not worsen colitis outcomes.

**Figure 7:**
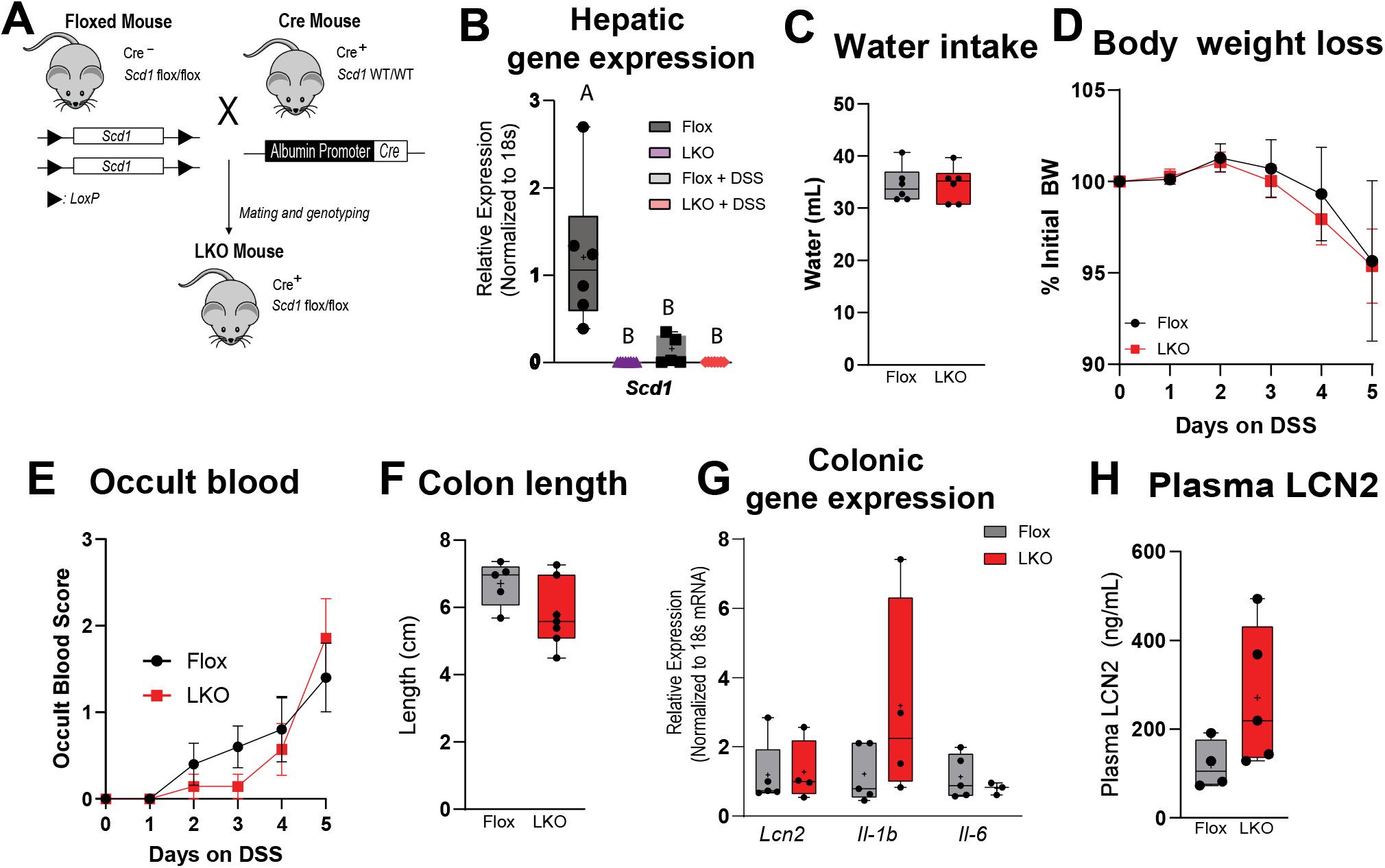
Mice lacking hepatic SCD1 are not more prone to DSS colitis. (A) Breeding strategy to generate LKO mice. (B) Gene expression measured via qRT-PCR. Mice were given 3% DSS in drinking water for 5 days and overall water intake (C) was measured. (D) Daily bodyweights and (E) daily occult blood scores were recorded. Occult blood scored as follows: 0, not occult blood; 1, positive occult blood; 2 visible blood in feces; 3, bloody anus. (F) Day 5 colon length was measured. (G) Colonic gene expression quantified via qRT-PCR and (H) plasma LCN2 measured via ELISA assay. n= 4-6 per group. Box plots show minimum and maximum values, + symbol at mean.

## Discussion

Consistent with prior reports [17, 23], we observed a significant reduction in SCD1 expression in the livers of both male and female mice upon treatment with a 5-day oral DSS regimen. This reduction occurs independently from modulation of *Srebp1c* expression in the liver. Interestingly, while a previous report indicated similar regulation of SCD1 in two other models of increased inflammation, i.e. a C*itrobacter rodentium* induced colitis model as well as upon direct intraperitoneal injection of LPS [23], we found that hepatic SCD1 was not suppressed in the *IL-10*^−/−^ model of chronic colitis. Concomitantly, mice on a chronic DSS regimen showed a reduction in hepatic SCD1, accompanied by a reduction in lipogenic transcription factor *Srepb-1c* and other downstream targe genes [17]. These data indicate important differences in hepatic responses to acute colonic inflammation as seen in the DSS, Citrobacter, and LPS models, vs. chronic DSS colitis and immune-mediated colitis in the *IL-10*^−/−^ mouse. Overall, the presence of colonic inflammation alone does not suppress hepatic SCD1 but rather, the downregulation of hepatic SCD1 is a response to acute colonic injury and inflammation. Interestingly, while the prior reports demonstrating Citrobacter- and LPS-mediated downregulation of hepatic SCD1 suggested a role for bacterial products in mediating this effect, we found similar downregulation of hepatic SCD1 by DSS administration even when using a model of antibiotic-mediated ablation. These results suggest that an intact gut microbiome is not essential for mediating the downregulation of hepatic SCD1 upon DSS treatment. While the precise mechanism by which acute DSS treatment reduces hepatic SCD1 expression remains elusive, it appears to be mediated in a manner independent of SREBP-1c and the gut microbiome.

It is intriguing that this specific downregulation of SCD1 alone, without coordinated changes in other lipogenic genes, is limited to the liver. In contrast, the adipose tissue response to acute DSS treatment includes not only a downregulation of SCD1 but also of most lipogenic genes, reflective of the catabolic state of the animal following this chemical insult. Previous studies have shown that longer term chronic DSS treatment ultimately results in downregulation of other lipogenic genes in the liver, as well [17]. However, in the short term, the primary hepatic changes in response to DSS include the significant reduction in SCD1 levels and concomitant lipid changes, as well as significant reductions in bile acid synthesis and altered cholesterol metabolism.

SCD1 catalyzes the conversion of SFAs to MUFAs, and it is well documented that tissue-specific inhibition of SCD1 leads to changes in lipidome composition [20–22]. Liver-specific SCD1-knockout mice show a reduction in hepatic monounsaturated fatty acids (MUFAs), specifically palmitoleate (16:1) and oleate (18:1) [20]. This observation was mirrored in skin-specific SCD1-KO mice, where skin total TAGs and wax di-esters were severely reduced [22]. Deletion of intestinal SCD1 results in decreased intestinal TAGs, DAGs and CE content, and desaturation ratios were reduced in the gut [21, 29]. Conversely, deletion of SCD2 in the gut results in upregulation of intestinal SCD1 and consequent increases in tissue desaturation ratios [30]. Thus, given the large reduction in hepatic SCD1 in DSS treated mice, we expected a significant reduction in liver TAGs and CEs in these mice. DSS treated mice did not show any changes in overall hepatic TAG content; however, MUFA-containing TAGs were significantly reduced (Figs. 3F, H). Additionally, we observed an increase in hepatic cholesterol esters, with unexpectedly increased desaturation ratios. This increased CE in the liver is likely an effect of reduced BA synthesis, as seen here (Fig. 4C) and reported by others [28]. Interestingly, it has been reported that intestinal dysbiosis in the mouse can lead to liver injury via activation of the NLRP3 inflammasome [31]. Links between inflammatory bowel disease and liver dysfunction have also been reported in human studies. A study showed a correlation between bile acid metabolism dysfunction and colitis where up to 80% of patients suffering from primary sclerosing cholangitis (PSC), chronic scaring of the bile duct causing bile buildup in the liver, also suffer from IBD [32]. Further, consistent with our observations of elevated CEs in livers of DSS-treated mice, a higher prevalence of metabolic-associated fatty liver disease (MAFLD) has been reported in patients with IBD [5, 33], with a NAFLD prevalence of 10-40% in IBD patients, depending on ethnical background [34, 35], all supporting a connection between colonic inflammation and altered hepatic lipid metabolism.

In addition to hepatic lipids, we also observed an increase in plasma lipids following DSS treatment, driven by an increase in SFAs and PUFAs. This resulted in decreased MUFA content of plasma fatty acids in DSS-treated mice. Increased plasma lipids are a known risk factor for the development of cardiovascular diseases (CVD). Multiple studies have shown that patients with inflammatory bowel diseases such as colitis and Crohn’s disease, have increased risk for developing CVD [36–39]. Previous studies investigating the plasma lipidomes of very high cardiovascular risk (VHCVR) patients have reported that these patients had increased levels of plasma glycerophospholipids [40] including significant increases in PC and PE, although no specific mechanisms were investigated. Another study examining plasma lipids in patients with coronary artery disease (CAD) reported that elevated plasma PE was associated with increased risk for CAD [41]. Interestingly, we found that these PL species were elevated in DSS-treated mice (Fig. 5D, Supplementary Fig. 2). Phospholipids are essential for maintaining the integrity of the colonic cell structure, with PC and PE, as well as PI and PS representing major phospholipids in colonocytes cell membrane [42]. Under conditions of colonic injury, such as DSS treatment, it is possible that more active remodeling of the colon, necessitated by damage to the gut barrier, may result in greater mobilization of PLs from the periphery and into circulation. Previous studies examining colonic mucosal lipids between healthy patients and active colitis patients showed major changes in glycerophospholipids, including PE, PC, PI, and PS, where many of these lipids were upregulated in colitis patients [43]. While this increase in plasma PLs may be a mechanism to limit and repair injury to the colon, chronic elevations in plasma PLs in colitis patients may also be linked to the elevated CAD risk observed in these patients.

Previous studies using models of whole-body deletion of SCD1 have provided contrasting results regarding the role of SCD1 in modulating colitis sensitivity. For instance, a report by Chen et al. indicated that *Scd1*−/−mice were more prone to DSS-colitis and that dietary oleate and adenoviral overexpression of hepatic SCD1 both protected WT mice from colitis [23]. *Scd1−/−* mice possess significant cutaneous abnormalities, leading to increase in trans-epidermal water loss and elevated water intake in this mouse strain [44, 45], which may have confounded the previous report which did not account for DSS dosage differences between genotype. To this effect, a subsequent study by Macdonald et al also noted that *Scd1−/−* mice consume more water than WT animals and that controlling DSS dosage in *Scd1−/−* animals by adjusting their water intake to match that of WT mice resulted in no greater sensitivity to colitis in *Scd1−/−* mice [46]. Our studies indicate that liver SCD1 expression is indeed downregulated by DSS treatment and that this downregulation is unique to SCD1 among the hepatic lipogenic genes. Our data also demonstrate, for the first time, that loss of hepatic SCD1 alone does not influence susceptibility to ulcerative colitis. Thus, while hepatic SCD1 overexpression or dietary oleate supplementation may ameliorate colitis as previously reported [23], the loss of liver SCD1 alone is insufficient to heighten colitis susceptibility.

In conclusion, our studies demonstrate that downregulation of hepatic SCD1 independently of SREBP-1c levels following acute colitis is not reflected in other tissues and is not causative of colitis severity. We also observe a reduction in total hepatic lipids, accompanied by an increase in liver CE and a severe dysregulation of cholesterol and bile acid metabolism following acute DSS treatment. Systemic lipid changes are also observed through increased plasma phospholipids and cholesterol, overall supporting a correlation between colonic inflammation and predisposition to CVD and liver dysfunction.

## Acknowledgments

We thank Dr. Karen Edelblum and Dr. Natasha Golovchenko for their help with colonic histology techniques. This work was funded by American Heart Association grant 20CDA35310305, NIH grant R01DK126963, and Crohn’s and Colitis Foundation of America grant 1450905 to HS, Rutgers Center for Lipid Research (RCLR) Small Research Grant to CD.

## Methods

### Colitis studies

Male and female mice were maintained on a 12-hour light/dark cycle with *ad libitum* access to water and a standard chow diet (Picolab 5053, LabDiet, NJ). 8–10-week-old mice (Jackson Laboratories) were individually housed 1-week prior to the start of the study. Mice were separated into a control group, receiving autoclaved water only, or a colitis group receiving 3% Dextran Sulfate Sodium (DSS), colitis grade (36,000-50,000, MP Biomedicals) in autoclaved water. Mice were treated for 5 days, and water/DSS was changed on day 3. Body weights were measured and feces collected daily to assess occult blood using Hemoccult guaiac fecal occult blood test (Beckman Coulter, Indianapolis, IN). On day 5, mice were fasted for 4 hours before being euthanized by isoflurane anesthesia followed by cardiac exsanguination. Plasma, livers, epididymal white adipose tissue (eWAT), and colons were collected. Prior to collection, colon tissue was flushed with ice-cold phosphate buffered saline (PBS), and connective tissue and fat was dissected off the tissue. Colon was cut into three equal parts denoted as “colon proximal” (proximal to cecum), “colon mid” and “colon distal”. All collected tissues were snap-frozen in liquid nitrogen and stored at −80°C or fixed in formalin for histology studies. For all *in vivo* procedures, every effort was made to minimize discomfort and suffering, in accordance with the protocols approved by the Animal Care and Use Committee of Rutgers University, New Brunswick, New Jersey.

### Gut microbiome ablation

Gut microbiome ablation was achieved via antibiotic (Abx) treatment, as previously described [47]. 8-week-old female mice were given drinking water containing a cocktail of antibiotics including ampicillin (1 g/L), neomycin (1 g/L), metronidazole (1 g/L) (all from Sigma, St. Louis, MO, United States) and vancomycin (VWR Radnor, PA, United States) (0.5 g/L) in autoclaved water for 2 weeks with daily monitoring. Following the 2-week ablation period, mice were randomly assigned to Water (control group) or DSS groups; mice continued to receive the Abx cocktail during the 5-days of water or DSS treatment. Fresh fecal pellets were collected before and after 2 weeks of Abx treatment and stored at −80°C until further analysis. Bacterial ablation was confirmed by isolating fecal bacterial DNA using the QIAamp DNA stool mini kit (QIAGEN, Germantown, MD, United States) following manufacturer’s instructions. Total fecal 16S rRNA was quantified by qPCR using forward primer 785F: 5′-GGA TTA GAT ACC CTG GTA and reverse primer 907R: 5′-CCG TCA ATT CCT TTR AGT TT, spanning the V5 region of 16S rRNA gene [48]. A 16S rRNA plasmid DNA with known copy number was used to generate a standard curve to quantify bacterial DNA in samples.

### Generation of IL-10−/−and liver Scd1^−/−^ mouse

BALB/cAnNTac (WT control for IL10 KO), and IL10 KO (BALB/BALB/cAnNTac-IL10em7Tac) mice were purchased from Taconic Biosciences. All mice were housed in a standard rodent facility and maintained on a commercial, irradiated chow diet (LabDiet® 5053, LabDiet, St. Louis, MO). Mice 16 weeks of age were used.

The strategy for generation of liver-specific SCD1 knockout mice has been previously described [20]. Briefly, C57BL/6 mice carrying flanking LoxP sequences surrounding exon 3 of the *Scd1* gene (*Scd1*^*lox/lox*^) were crossed with C57BL/6 mice expressing Cre-recombinase under the albumin promoter to generate *Scd1*^*lox/lox; Cre+*^ (LKO) mice. Littermates not expressing Cre (*Scd1*^*lox/lox*^) were used as the control (floxed) group. Mice were maintained on a 12-hour light/dark cycle with *ad libitum* access to water and standard chow diet (Picolab 5053, LabDiet, NJ).

### Histology

Colons were flushed with ice-cold PBS, cut longitudinally to expose the lumen, and rolled in a “Swiss-roll” prior to fixation in 10% formalin for 24h, followed by paraffin embedding. Tissue slices mounted on charged slides were stained with hematoxylin and eosin for structural visualization. Images were visualized using a light microscope (Echo Revolve).

### Gene expression analyses

RNA was isolated using RNAzol RT (Cat # RN190, Molecular Research Center, Ohio, USA) or RNeasy Lipid Tissue kit (eWATQiagen, Germantown, MD), following manufacturer’s instructions. Due to the nature of DSS, which can inhibit reverse transcriptase, samples were additionally purified with lithium chloride (LiCl) with small modifications to a previously published protocol [49]. Briefly, equal volume of LiCl Precipitation Solution (Cat # AM9480, Invitrogen by Thermo Fisher Scientific, Vilnius, Lithuania) was added to the volume of resuspended RNA pellet, vortexed and stored at −20°C overnight. Samples were then centrifuged at 4°C, maximum speed, for 20min.

Pellets were washed twice with 75% ethanol before being resuspended in RNAse-free water. RNA concentration was quantified using a NanoDrop 2000 spectrophotometer (Thermo Scientific, Waltham, MA, United States). 1 μg of RNA was used to synthesize complementary DNA (cDNA) using Superscript III first-strand synthesis system (Invitrogen by Thermo Fisher Scientific, Waltham, MA, United States). Quantitative real-time PCR (qRT-PCR) was performed on a QuantStudio 3 Real-Time PCR System (Applied Biosystems) with gene-specific primers (table 1). Data were normalized to the expression of *18s rRNA* for liver and WAT tissues, and to *Gusb* for colon samples. Quantification was carried out using the 2^-∆ ∆Ct^ method [50]. Primer sequences found in table 1.

### Protein analyses

Whole cell lysates were prepared, and protein concentrations were measured using the Pierce BCA protein assay (ThermoFisher Scientific, Waltham, MA). Equal amounts of protein were separated via SDS-PAGE and transferred to PVDF membranes. Total protein staining was performed using Ponceau stain (Aventor, Radnor, PA) to confirm uniform protein loading r. Membranes were blocked in 3% milk in Tris-buffered saline (TBS, 150nM NaCl, 50mM Tris-HCl, pH 7.4) with 0.1% Tween 20 for 1 hour at room temperature with rocking. Primary antibodies were diluted 1:1000 in 3% milk in TBS-Tween 20. Membranes were incubated with primary antibodies, followed by incubation with HRP- or Alexa-fluor conjugated secondary antibodies, and signal was detected using enhanced chemiluminescence or fluorescence imaging, respectively, on an Azure c600 imaging system (Azure Biosystems, Dublin, CA, USA). Primary antibodies used in this study were as follows: SCD1 (C12H5, Cell Signaling Technology, Danvers, MA, USA), GAPDH (D16H11, Cell Signaling Technology, Danvers, MA, USA), β-actin (A5316, Millipore Sigma, Burlington, MA, USA).

### Liver Lipidomics

Hepatic lipids were extracted by a modified Folch method, separated by thin-layer chromatography, and analyzed by GC-MS as previously described [21, 51–53]. MassHunter Data Acquisition software and MassHunter Quantitative Analysis software were utilized for peak analyses. Peak identity was confirmed by National Institute of Standards and Technology (NIST) library search. Fatty acids were quantified using specific standard curves generated for each acyl chain.

### Plasma lipidomics

Acyl chains composition of total plasma lipids was analyzed by GC-MS, as previously described [21, 51–53]. In addition, unbiased Plasma lipidomic profiling was performed using Ultra Performance Liquid Chromatography-Tandem Mass Spectrometry (UPLC-MS/MS) by the Columbia University Lipidomics Core, as has been previously reported [54, 55]. Lipid species were quantified using multiple reaction monitoring (MRM) transitions [56–58] under both positive and negative ionization modes in conjunction with referencing of appropriate internal standards. Lipid levels for each sample were calculated by summing the total number of moles of all lipid species measured by all three LC-MS methodologies and then normalizing the total to mol %. Total lipid abbreviations are as follows: (FC) Free Cholesterol, (CE) Cholesterol Ester, (AC) Acyl Carnitine, (MG) Monoacylglycerol, (DG) Diacylglycerol, (TAG) Triacylglycerol, (Cer) Ceramide, (dhCer) Dihydroceramide, (SM) Sphingomyelin, (dhSM) Dihydrosphingomyelin, (Sulf) Sulfatide, (MHCer) Monohexosylceramide (galactosylceramide + glucosylceramide), (LacCer) Lactosylceramide, (GM3) Monosialodihexosylganglioside, (GB3) Globotriaosylceramide, (PA) Phosphatidic acid, (PC) Phosphatidylcholine, (PCe) Alkyl phosphatidylcholine, (PE) Phosphatidylethanolamine, (PEp) Plasmalogen phosphatidylethanolamine, (PS) Phosphatidylserine, (PI) Phosphatidylinositol, (PG) Phosphatidylglycerol, (BMP) Bis(monoacylglycero)phosphate, (AcylPG) Acyl Phosphatidylglycerol, (LPC) Lysophosphatidylcholine, (LPCe) Alkyl lysophosphatidylcholine, (LPE) Lysophosphatidylethanolamine, (LPEp) Plasmogen lysophosphatidylethanolamine, (LPI) Lysophosphatidylinositol, (LPS) Lysophosphatidylserine, (NAPE) N-Acyl Phosphatidylethanolamine, (NAPS) N-Acyl Phosphatidylserine, (NSer) N-Acyl Serine

### Plasma lipoprotein fractionation and cholesterol assessment

Lipoprotein fractions from plasma of control and DSS-treated mice were isolated by density gradient ultracentrifugation, as previously described [59, 60]. Briefly, 400μL of potassium bromide (KBr) (d = 1.0063 g/mL) was floated over 100μL of plasma in a 0.5mL microcentrifuge tube (Beckman Coulter, Cat # 343776). After centrifugation at 140,000 x g at 10°C for 2.5 hours, the top layer containing the VLDL + chylomicrons fraction (d < 1.006 g/mL; exactly 100 μL) was removed with a Hamilton glass syringe (Hamilton Co.) and saved. The remaining solution was brought to 500μL with KBr for a final density of 1.063g/mL and centrifuged at 160,000 x g at 10°C overnight. The top 100 μL containing LDL (d < 1.06 g/mL) was collected and saved. The remaining solution was brought up to 500μL with KBr for a final density of 1.21 and centrifuged at 160,000 x g at 10°C overnight. The top 100μL containing the HDL fraction (d = 1.063–1.21 g/mL) was collected and saved. All collected lipoprotein fractions were stored at −80°C until further analysis.

Total plasma cholesterol was measured using the Cholesterol E kit colorimetric assay (Fujifilm, Cat # 999-02601) following manufacturer’s directions.

### Plasma LCN2 measurement

Plasma Lipocalin2 (LCN2) was measured via DuoSet enzyme-linked immunosorbent assay (ELISA) kit (Cat # DY1857) from R&D Systems (Minneapolis, MN, United States), following manufacturer’s directions.

### Statistical analysis

Data are expressed as mean ± standard error of the mean (SEM) for biological replicates or in box plots with mean, minimum, and maximum Comparisons are carried out using the Student *t-* test for 2-group comparisons or one-way ANOVA followed by post-hoc analysis using a multiple comparison procedure with Bonferroni/Dunn post-hoc comparison in Graph Pad Prism (version 10.5.0 for windows, GraphPad Software, La Jolla, CA, USA). *p* values <0.05 were considered statistically significant.

