## Supplementary Figure 1 for "Hepatic stearoyl-CoA desaturase-1 is specifically suppressed by dextran sodium sulfate but does not influence colitis sensitivity"

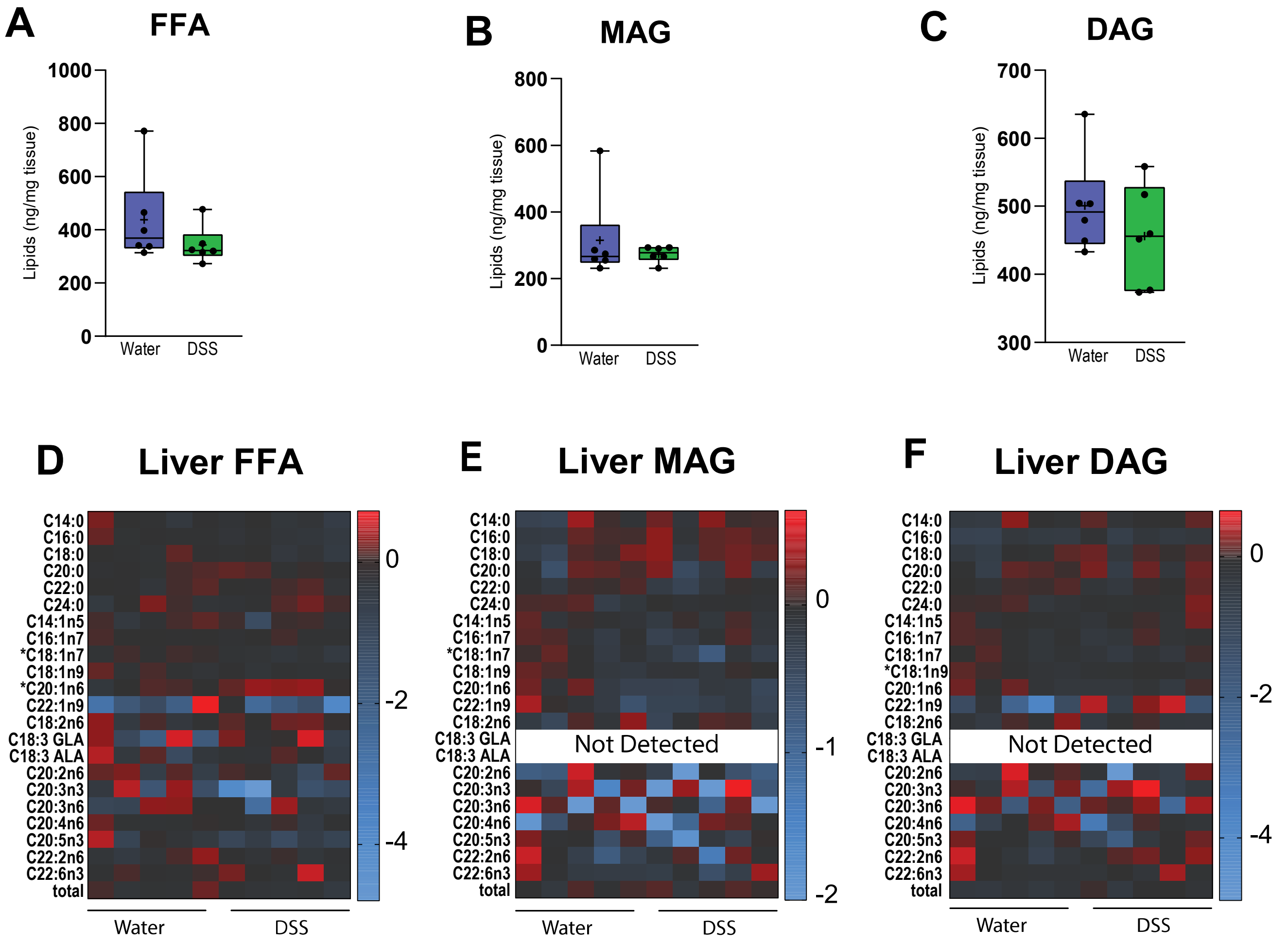


**Supplementary figure 1: Liver lipid composition of FFA, MAG and DAG following DSS.** (A-F) Total content of free fatty acids (FFA), monoacylglycerols (MAG) and diacylglycerols (DAG) was determined by TLC-GC-MS using liver tissue. n=6 per group. *p<0.05 vs water group.
