## Supplementary Figure 2 for "Hepatic stearoyl-CoA desaturase-1 is specifically suppressed by dextran sodium sulfate but does not influence colitis sensitivity"

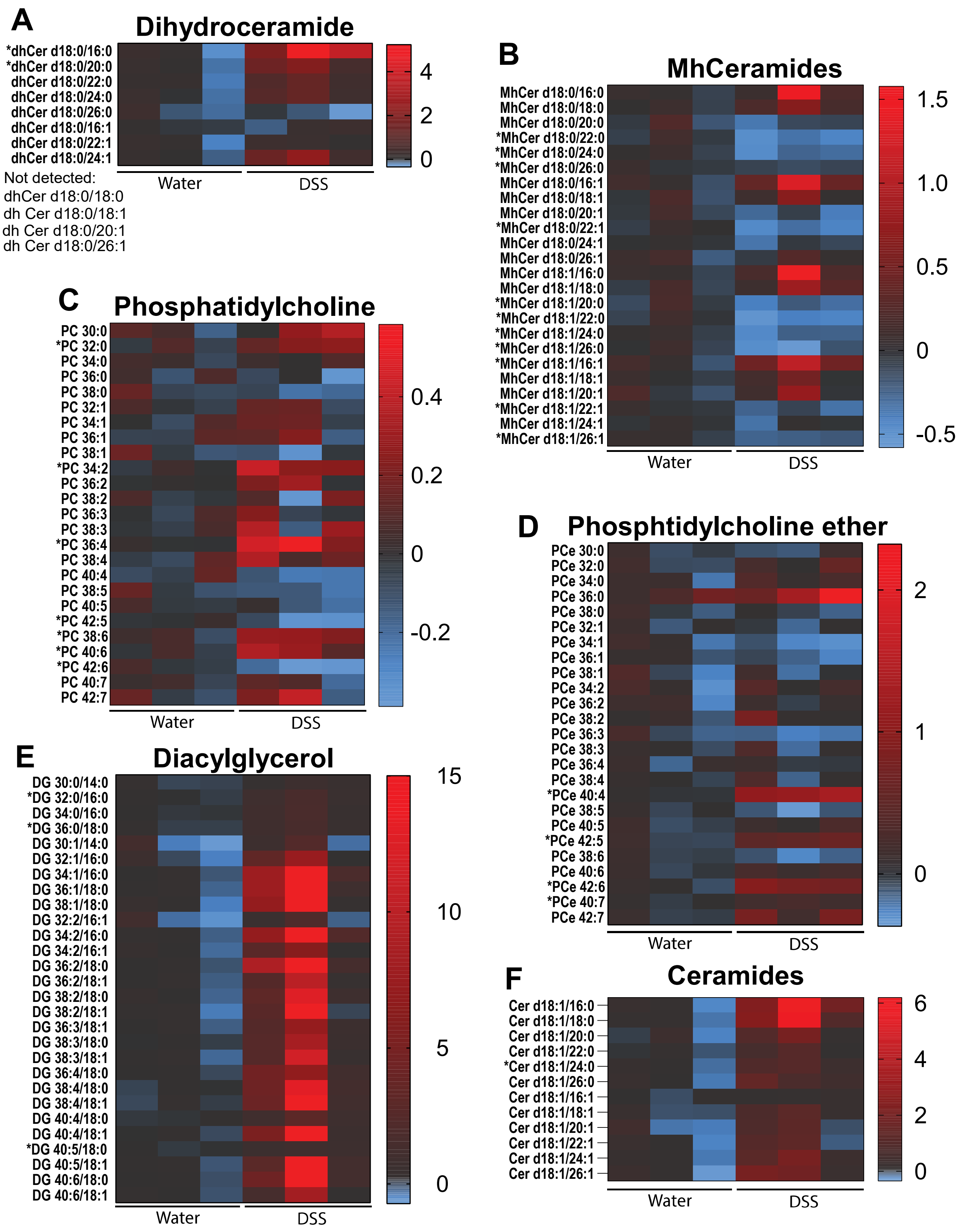

A
Dihydroceramide
*dhCer d18:0/16:0
*dhCer d18:0/20:0
dhCer d18:0/22:0
dhCer d18:0/24:0
dhCer d18:0/26:0
dhCer d18:0/16:1
dhCer d18:0/22:1
dhCer d18:0/24:1
Not detected:
dhCer d18:0/18:0
dh Cer d18:0/18:1
dh Cer d18:0/20:1
dh Cer d18:0/26:1
Water
DSS
4
2
0
B
MhCeramides
MhCer d18:0/16:0
MhCer d18:0/18:0
MhCer d18:0/20:0
*MhCer d18:0/22:0
*MhCer d18:0/24:0
*MhCer d18:0/26:0
MhCer d18:0/16:1
MhCer d18:0/18:1
MhCer d18:0/20:1
*MhCer d18:0/22:1
MhCer d18:0/24:1
MhCer d18:0/26:1
MhCer d18:1/16:0
MhCer d18:1/18:0
*MhCer d18:1/20:0
*MhCer d18:1/22:0
*MhCer d18:1/24:0
*MhCer d18:1/26:0
*MhCer d18:1/16:1
MhCer d18:1/18:1
MhCer d18:1/20:1
*MhCer d18:1/22:1
MhCer d18:1/24:1
*MhCer d18:1/26:1
Water
DSS
1.5
1.0
0.5
0
-0.5
C
Phosphatidylcholine
PC 30:0
*PC 32:0
PC 34:0
PC 36:0
PC 38:0
PC 32:1
PC 34:1
PC 36:1
PC 38:1
*PC 34:2
PC 36:2
PC 38:2
PC 36:3
PC 38:3
*PC 36:4
PC 38:4
PC 40:4
PC 38:5
PC 40:5
*PC 42:5
*PC 38:6
*PC 40:6
*PC 42:6
PC 40:7
PC 42:7
Water
DSS
0.4
0.2
0
-0.2
D
Phosphtidylcholine ether
PCe 30:0
PCe 32:0
PCe 34:0
PCe 36:0
PCe 38:0
PCe 32:1
PCe 34:1
PCe 36:1
PCe 38:1
PCe 34:2
PCe 36:2
PCe 38:2
PCe 36:3
PCe 38:3
PCe 36:4
PCe 38:4
*PCe 40:4
PCe 38:5
PCe 40:5
*PCe 42:5
PCe 38:6
PCe 40:6
*PCe 42:6
*PCe 40:7
PCe 42:7
Water
DSS
2
1
0
E
Diacylglycerol
DG 30:0/14:0
*DG 32:0/16:0
DG 34:0/16:0
*DG 36:0/18:0
DG 30:1/14:0
DG 32:1/16:0
DG 34:1/16:0
DG 36:1/18:0
DG 38:1/18:0
DG 32:2/16:1
DG 34:2/16:0
DG 34:2/16:1
DG 36:2/18:0
DG 36:2/18:1
DG 38:2/18:0
DG 38:2/18:1
DG 36:3/18:1
DG 38:3/18:0
DG 38:3/18:1
DG 36:4/18:0
DG 38:4/18:0
DG 38:4/18:1
DG 40:4/18:0
DG 40:4/18:1
*DG 40:5/18:0
DG 40:5/18:1
DG 40:6/18:0
DG 40:6/18:1
Water
DSS
15
10
5
0
F
Ceramides
Cer d18:1/16:0
Cer d18:1/18:0
Cer d18:1/20:0
Cer d18:1/22:0
*Cer d18:1/24:0
Cer d18:1/26:0
Cer d18:1/16:1
Cer d18:1/18:1
Cer d18:1/20:1
Cer d18:1/22:1
Cer d18:1/24:1
Cer d18:1/26:1
Water
DSS
6
4
2
0

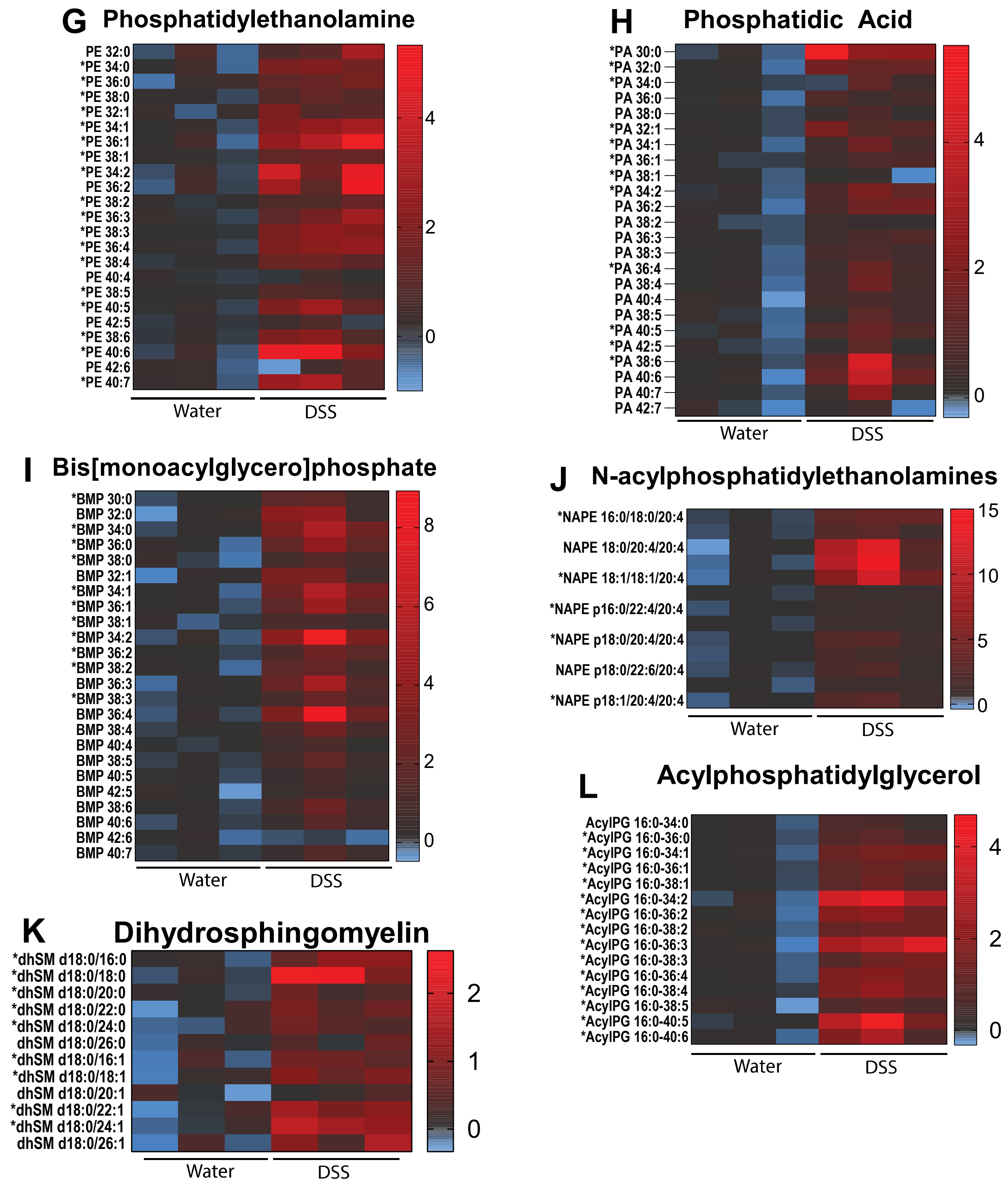


**Supplementary figure 2: Plasma lipidomics following DSS treatment**. (A-L) Plasma lipidomic content determined via UPLC-MS/MS. n=3 per group.
