## Supplementary material for "Hepatic stearoyl-CoA desaturase-1 is specifically suppressed by dextran sodium sulfate but does not influence colitis sensitivity": Table 1

| Gene Name | Forward Primer (5' to 3') | Reverse Primer (3' to 5') |
|---|---|---|
| *18s-rRNA* | TCACCATCATGCAGAACCCA | CCTGGCTGTACTTCCCATCCT |
| *Abca1* | GGACATGCACAAGGTCCTGA | CAGAAAATCCTGGAGCTTCAAA |
| *Acat1* | GGAAGTTGGGTGCCACTTCG | GGTGCTCTCAGATCTTTGG |
| *Acat2* | GACTTGGTGCAATGGACTCG | GGTCTTGCTTGTAGAATCTGG |
| *Acc* | TGACAGACTGATCGCAGAGAAAG | TGGAGAGCCCCACACACA |
| *Atgl* | CTTGAGCAGCTAGAACAATG | GGACACCTCAATAATGTTGGC |
| *Bsep* | TGAATGGACTGTCGGTATCTGTG | CCACTGCTCCCAACGAATG |
| *Cd36* | TACGCTGTGTTCGGATCTGA | ATTCTGGAGGGGTGATGCAA |
| *Cpt1a* | TGAGTGGCGTCCTCTTTGG | TCAGCGAGTAGCGCATAGTCA |
| *Cyp27a1* | GCCTCACCTATGGGATCTTCA | TCAAAGCCTGACGCAGATG |
| *Cyp7a1* | AACAACCTGCCAGTACTAGATAGC | GTGTAGAGTGAAGTCCTCCTTAGC |
| *Dgat1* | CTGGATTGTGGGCCGATTCT | ATACATGAGCACAGCCACCG |
| *Dgat2* | GCCTGCAGTGTCATCCTCAT | TGGGCGTGTTCCAGTCAAAT |
| *Fas* | GCTGCGGAAACTTCAGGAAAT | AGAGACGTGTCACTCCTGGACTT |
| *Fxr* | TCCGGACATTCAACCATCAC | TCACTGCACATCCCAGATCTC |
| *Gusb* | CCGATTATCCAGAGCGAGTATG | CTCAGCGGTGACTGGTTCG |
| *Hmgr* | CTTGTGGAATGCCTTGTGATT | AGCCGAAGCAGCACATGAT |
| *Il-1b* | TTGACGGACCCCAAAAGATG | AGAAGGTGCTCATGTCCTCAT |
| *Il-6* | ATGAACAACGATGATGCACTT | TATCCAGTTTGGTAGCATCCAT |
| *Lcad* | GAAAGGCTCTTAATTGCTGAG | ATTCTGCTAGTTTATGCTGC |
| *Lcn2* | TGCCACTCCATCTTTCCTGTT | GGGAGTGCTGGCCAAATAAG |
| *Ldlr* | AGGCTGTGGGCTCCATAGG | TGCGGTCCAGGGTCATCT |
| *Mcad* | GCTAGTGGAGCACCAAGGAG | CCAGGCTGCTCTCTGGTAAC |
| *Mrp2* | CTGAGTGCTTGGACCAGTGA | GTTAACAGCTGCCTGTGCAA |
| *Mrp3* | TGAGATCGTCATTGATGGGC | AGCTGAGAGCGCAGGTCG |
| *Mttp* | AACCGATTAACCTGGGTCAAGA | ACCGGCGACAACAGTGTTTA |
| *Ntcp* | GGCCACAGACACTGCGCT | AGTGAGCCTTGATCTTGCTGAACT |
| *Oatp1* | GGGAACATGCTTCGTGGGATA | GGAGTTATGCGGACACTTCTC |
| *Ost-β* | GTATTTTCGTGCAGAAGATGCG | TTTCTGTTTGCCAGGATGCTC |
| *Pgc-1a* | TCGATGTGTCGCCTTCTTGC | ACGAGAGCGCATCCTTTGG |
| *Ppar-a* | ACGATGCTGTCCTCCTTGATG | GTGTGATAAAGCCATTGCCGT |
| *Scarb1* | TCCCTCATCAAGCAGCAGGT | TTCCACATCCCGAAGGACA |
| *Scd1* | CCTCTGGAGCCACAGAACTT | GCCATGGTGTTGGCAATGAT |
| *Scd2* | CCACTTGAAAGTAGCCTTAC | ATAGAATAGGGCCACAGCTCA |
| *Sirt1* | GTCTCCTGTGGGATTCCTGA | CAAACATGGCTTGAGGGTCT |
| *Srebp-1c* | GGAGCCATGGATTGCACATT | GGCCCGGGAAGTCACTGT |

**Table 1 –** qRT-PCR primer sequences.
